# Is apparent digestibility associated with residual feed intake and enteric methane emission in Nellore cattle?

**DOI:** 10.1101/2022.12.01.518681

**Authors:** Sarah Bernardes Gianvecchio, Leandro Sannomiya Sakamoto, Luana Lelis Souza, Lorena Ferreira Benfica, Juliana de Oliveira Santos Marcatto, Eduardo Marostegan de Paula, Roberta Carilho Canesin, Joslaine dos Santos Gonçalves Cyrillo, Lucia Galvão Albuquerque, Maria Eugênia Zerlotti Mercadante

## Abstract

Residual feed intake (RFI) detects animals that consume less dry matter (DM) adjusted for production and maintenance, and differences in the digestibility may explain differences in RFI among contemporaneous cattle. The aim of the study was to evaluate the relationship between digestibility, RFI, and enteric methane emission in growing Nellore cattle (*Bos indicus*) divergently classified based on RFI phenotypes and also on RFI breeding value. One hundred twenty-two Nellore cattle submitted to performance testing in two test groups (249±4.33 days of age, and 332±4.22 days of age at the beginning of the test) were classified based on residual feed intake (RFI). A sample of 80 animals classified as low RFI (−0.748±0.076 kg DM/day) or high RFI (0.775±0.075 kg DM/day) was evaluated regarding feed compounds digestibility, fecal excretion, and enteric methane (CH_4_, g/day) emission. There was no significant difference in feed compounds digestibility between the most and least efficient animals. However, DM intake (6.92 vs. 8.66 kg DM/day) and feed conversion (7.93 vs. 9.42 kg/kg) were lower in low RFI animals. On average, low RFI animals emitted 14.3 g less CH_4_ per day (174 vs. 188 g CH_4_/day; P=0.02); however, CH_4_ emission expressed as g/kg DM intake (23.1 vs. 20.1; P<0.01) and the percentage of gross energy intake lost as CH_4_ (8.13 vs. 7.08%; P<0.01) were higher in these animals. There are clear benefits (economic and environmental) of using more feed efficient animals in the beef production chain, i.e., animals that exhibit lower feed intake, lower fecal excretion and lower enteric methane emission without differences in weight gain or body weight. Variations in feed efficiency among them cannot be explained by differences in DM or feed compounds digestibility. More efficient animals emit less enteric methane (CH4, g/day) than less efficient animals, probably as a result of lower DMI.

## Introduction

The growth of the world population and the global demand for food raise concerns regarding climate change since the increase in the emission of greenhouse gases from agricultural production systems is proportional to the increase in food production [1].

Residual feed intake (RFI) detects animals that consume less dry matter (DM) adjusted for weight gain and metabolic body weight, and may therefore be a potential measure to identify animals that emit less methane (CH_4_, g/day) without affecting performance [2–4]. Differences in the digestibility may explain the differences in RFI among contemporaneous cattle [5]. Studies have reported a negative correlation of RFI with DM digestibility [6–9], as well as with the digestibility of other feed compounds [10], i.e., more efficient animals in terms of reducing feed intake without changing the average daily gain exhibit greater digestibility. However, other studies failed to show this relationship [11–13], or even reported a positive correlation between digestibility and RFI [14]. It is therefore not clear whether the increase in the digestive capacity of efficient animals (low RFI) is inherent to the individual or is due to the slower passage of digesta through the rumen caused by lower dry matter intake [15] when compared to inefficient animals (high RFI).

The emission of methane resulting from the digestive process reduces the efficiency of energy utilization and performance of animals [16]. More efficient diet utilization is particularly important since it reduces feed intake and methane emission for the same production level [17], as well as fecal production [18].

The hypothesis of this study is that extremely efficient animals (low RFI) exhibit greater digestibility and consequently greater feed compounds utilization, associated with lower enteric methane emission, than extremely inefficient animals (the most efficient tercile compared to the least efficient tercile). The novelty of the present study, in relation to those reviewed by Kenny et al. [15] and Cantalapietra-Hijar et al. [19], is that it involves a considerable number of animals of indicine cattle fed a high forage diet. The aim of the present study was to evaluate the relationship between digestibility, RFI, and enteric methane emission in growing Nellore cattle (*Bos indicus*) divergently classified based on RFI phenotypes and also on RFI breeding value.

## Materials and Methods

The study was approved by the Ethics Committee on Animal Experimentation of the Institute of Animal Science (Protocol 86 Number 278–19), Brazil. The animals included in the study are from a selection experiment of Nellore cattle for growth traits established in 1980 at the Beef Cattle Research Center, Institute of Animal Science, Sertãozinho, SP, Brazil (21°10′ south latitude and 48°5′ west longitude), and since 2004 the animals are measured for feed efficiency traits after weaning [20]. Excepting during performance tests, animals were kept on *Brachiaria brizantha* (Urochloa spp) pasture.

### Performance test

A total of 122 non-castrated males born in 2018 were evaluated in two consecutive test groups (TG1 [18June to 09September2019], n: 60, mean: 249±3.59 days of age, and mean: 224±5.39 kg of live weight at the beginning of the test; TG2 [11September to 03December2019], n: 62, mean: 329±3.53 days of age, and mean: 285±5.30 kg of live weight at the beginning of the test). The performance tests had a duration of 83 days, preceded by 28 days for adaptation to the diet. The animals were weighed every 15 days without previous fasting, always before the first food delivery of the day. The animals were kept in a collective pen equipped with ten electronic feed bunks (GrowSafe^®^, Airdrie-AB, Canada) for the automatic recording of individual daily feed intake, with *ad libitum* access to diet and water. The daily intake records were discarded when the Growsafe System’s assigned feed disappearance metric was less than 90%.

The diet (Table 1) was formulated for an average daily gain of 1.100 kg/animal [21] and supplied twice a day (9:00 and 15:00 h). Samples of the ingredients were collected weekly (forage) and monthly (concentrate) for quantification of the DM content of the diet and chemical analysis.

**Table 1.**
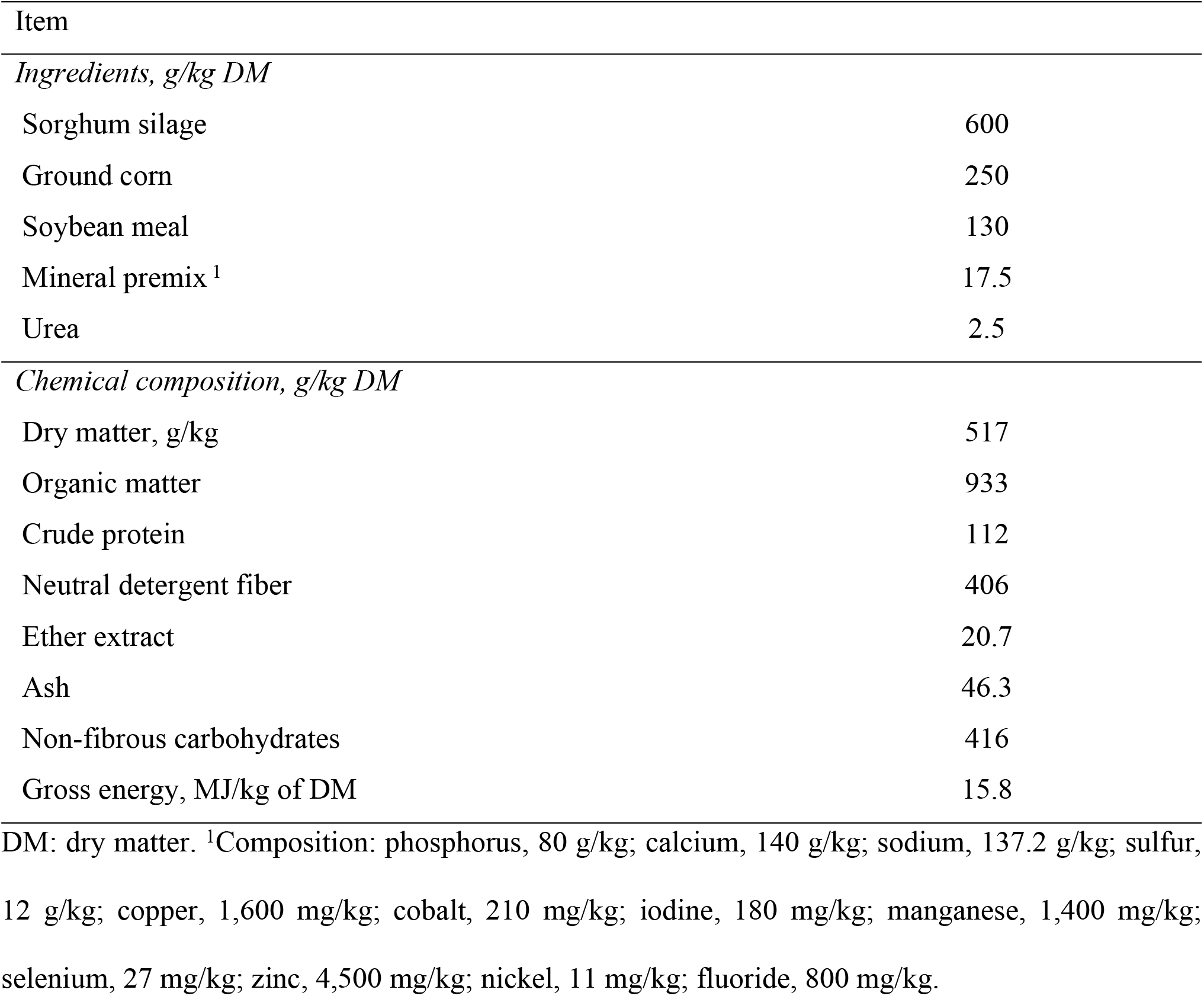
Ingredients and chemical composition of the diet

The dry matter intake (DMI) was obtained as the mean of all valid days (66 valid days for TG1 and for TG2) during the period. The average daily gain (ADG) was estimated by the linear regression coefficient of weights on days in test (DIT) according to the equation: yi = α + β × DITi + Ɛi, where yi is the weight of the animal in the i^th^ observation; α is the intercept representing the initial weight of the animal; β is the linear regression coefficient representing ADG; DITi is the day in test in the i^th^ observation; Ɛi is the random error associated with each observation. Estimate of the midpoint metabolic body weight (mBW^0.75^) was obtained as follows: mBW^0.75^ = (BWi + (0.5 DOT x ADG))^0.75^, where BWi is the initial body weight and DOT is the total of days on test. The RFI was calculated as the difference between observed and expected DMI, which was estimated by multiple regression of DMI on ADG and mBW^0.75^ within each test group using the GLM procedure (SAS Inst., Inc., Cary, NC). The equations of predicted DMI were: DMIp = 1.298 (± 1.107) + 2.706 (± 0.860) x ADG + 0.060 (±0.024) x mBW^0.75^ (R^2^ = 0.53) for TG1, and DMIp = −0.055 (± 0.719) + 2.151 (± 0.584) x ADG + 0.087 (±0.013) x mBW^0.75^ (R^2^ = 0.75) for TG2. Feed conversion ratio was obtained as the ratio between DMI and ADG.

At the end of the performance tests, the animals were classified based on RFI. Forty animals of each extreme tercile were used for the analysis of feed compounds digestibility and enteric methane emission: low RFI (mean: −0.774±0.068 kg DM/day; 17 animals from TG1 and 23 animals from TG2) and high RFI (mean: 0.782±0.068 kg DM/day; 22 animals from TG1 and 18 animals from TG2). Next, the breeding values for RFI (EBV-RFI) were estimated for these animals in a genetic evaluation of 1,878 animals with the RFI phenotype, 2,256 animals with genomic data, and 12,594 animals in the relationship matrix using the single-step GBLUP method, as described by Benfica et al.[20]. Since the herd has been selected for lower RFI and higher body weight since 2008, the EBV-RFI was expressed as a deviation of the average EBV-RFI of all animals born in 2008 (−0.0384 kg DM/day); and there were more negative EBV-RFI animals (n=56) than positive EBV-RFI animals (n=24).

### Estimates of digestibility

To estimate fecal excretion using indigestible neutral detergent fiber (iNDF) as the internal marker, diurnal feces samples were collected from the rectum of the animals for two consecutive days at defined times (7:30 and 13:30 h, and 10:30 and 16:30 h) [22] in the first (initial period) and last week (final period) of the performance test of each test group. The feces samples (300 g/animal) and dietary ingredients of the two fecal collection periods were stored at −20°C and then pre-dried in a forced ventilation oven at 60±5°C for 72 hours and ground in a Wiley mill (2-mm sieves). For the determination of iNDF, dietary ingredient and fecal samples were incubated (6 replicates per period + 1 blank) in four Nellore animals fed the same diet as used in the performance test by the *in situ* non-woven textile bag method (TNT - 100 g/m^2^, 4×5 cm at a proportion of 25 mg DM/cm^2^ surface area) for 264 hours, as described by Casali et al. [23]. The samples were then submitted to extraction in neutral detergent ([24], adapted to the TECNAL recommendations) using a fiber determinator (TE 149; TECNAL, Piracicaba, SP, Brazil).

Total daily fecal excretion of the animal (FE, g DM/day) was estimated as follows [25]: FE = D/[M], where *D* is the daily intake of the internal marker (g DM/day) of the percentage of dietary iNDF and DMI during fecal collection (DMIc), and *[M]* is the content of the marker in the fecal sample (g/g). The DMIc was obtained as the mean of all valid days of feed intake during the fecal collection period (4 days prior to collection, 2 days of collection, and 1 day post-collection per period). The FE was expressed as the mean of the initial and final periods. Apparent DM and feed compounds digestibility (D, g/kg) was obtained as follows [26]: D = [(consumed-excreted)/consumed], and was expressed as the mean apparent digestibility of the initial and final periods.

### Measurement of enteric methane emission

Enteric methane emission (CH_4_, g/day) was estimated using the sulfur hexafluoride (SF_6_) tracer gas technique, adapted to the local conditions, following the recommendations of Berndt et al. [27]. The animals were allowed to adapt for 10 days to the sampling apparatus, leather halter, and light saddle. About 250 permeation tubes (brass capsule) containing SF_6_ were maintained in an oven at 39°C and calibrated by weekly weighing on an analytical scale for 8 weeks before the beginning of sampling in order to ensure constant and linear emission of SF_6_. Eighty capsules with an average release of 4.598±0.040 and 2.460±0.039 mg SF_6_/day in the first (TG1) and second (TG2) samplings, respectively, were selected and administered 5 days before the beginning of the gas sampling period.

Starting between 7:30 and 8:00 h, methane was collected on consecutive days [24June to 02July2019 for TG1; 15 to 23October2019 for TG2], with exchange of the cylinder every 24 hours in order to ensure five samples per animal. The period was extended up to 8 days due to losses. The gases expelled from the mouth and nostrils of the animals were captured in a controlled and continuous manner through a stainless steel capillary tube, which was calibrated as described by Deighton et al. [28] and protected by a flexible hose fixed to the halter and connected to the collection cylinder. The latter was subjected to vacuum (0 atm) and attached to the animal’s back. The pressures (initial and final) of the cylinders were monitored daily to ensure the quality of the sample. The concentrations of environmental gases were obtained by daily samplings at strategical points in the paddock, collected into two cylinders. At the end of each sampling period, the sampled gases were analyzed by gas chromatography (HP6890, Agilent, Wilmington, DE, USA) using a flame ionization detector for CH_4_ and an electron capture detector for SF_6_. The amount of enteric methane was estimated as a function of SF6 concentrations, relating the results to the known rate of tracer gas release by the capsule deposited in the rumen, with correction for sampled environmental concentrations and molecular weights [27].

The daily methane emission (CH_4_, g/day) of each animal is the arithmetic mean of emissions estimated in the five samples. Methane emission was also obtained in relation to midpoint body weight (CH_4_/mBW, g/kg/day), average daily gain (CH_4_/ADG, g/kg/day), and dry matter intake (CH_4_/DMI, g/kg/day), as well as percentage (%) of gross energy intake lost in the form of methane.

### Chemical analyses (diet and feces)

The diet and feces samples were ground to pass a 1-mm screen (Wiley mill) and analyzed for DM (method 934.01), ash, and organic matter (method 938.08), and ether extract (method 954.02) content according to the AOAC [29]. Total nitrogen (total N x 6.25 = crude protein) was analyzed using a combustion assay (DUMATHERM^®^ N Analyzer, Gehart, Germany) according to AOAC International [30] method 990.13. For neutral detergent fiber (NDF), samples were treated with alpha thermo-stable amylase omitting sodium sulfite determined according to the method of Van Soest et al. [31], and adapted for fiber determinator (TE-149; TECNAL, Piracicaba, 187 SP, Brazil).

Non-fibrous carbohydrates (NFC) were obtained by NFC = 100-(%CP+%EE+%ash+%NDF). The energy value of the diet was estimated from the results of dietary chemical analysis: TDN = %CPD+%NFCD+%NDFD+2.25×%EED, in which *D* is the digestibility (NRC, 2000). Gross energy was determined with an adiabatic calorimeter (IKA WERKE Model C5003) according to the method of Parr [32]. Digestible energy of the diet was calculated by subtracting total fecal gross energy from total dietary gross energy.

### Statistical analysis

The variables were analyzed using the MIXED procedure (SAS Inst., Inc., Cary, NC), fitting a model that included the fixed effect of RFI class (i=1, 2), and the linear effect of initial age of the animal as covariate, and the test group (j=1, 2) as random effects. Means were adjusted by the least-squares method (LSMEANS). Complementary analyses were performed that included the fixed effect of EBV-RFI (i=1, 2). The quadratic effect of initial age was included when the linear effect was significant (P<0.05). The fixed interactions between RFI class (or EBV-RFI class) and test group class were tested and included in the final models when significant (P<0.05). Different variances residuals for the test group (i=1, 2) were modeled using the GROUP option of the REPEATED command for all variables studied.

The phenotypic correlations of CH_4_ (g/day) and CH_4_/DMI (g/kg/day) with apparent DM and feed compounds digestibility were estimated using the CORR procedure (SAS Inst., Inc., Cary, NC). Statistical significance was declared when P<0.05.

## Results

Table 2 shows the performance, feed efficiency, and estimated fecal excretion, fecal excretion and apparent digestibility. There was no significant difference in apparent digestibility between most and least efficient animals. Dry matter intake (6.92 vs. 8.66 kg DM/day) and feed conversion (7.93 vs. 9.42 kg/kg) were −20.1% and −15.8%, respectively, in low RFI animals compared to their high RFI counterparts; consequently, estimated fecal excretion were also lower in low RFI animals. There was no difference in the performance of low and high RFI animals. As a complementary analysis, negative EBV-RFI and positive EBV-RFI classes were compared, confirming the results obtained for the comparison of extreme RFI phenotype classes in terms of performance, excretion, and digestibility. The rank correlation between RFI phenotype and EBV-RFI of these 80 animals was high (r=0.79).

**Table 2.** Performance, feed efficiency, and estimated fecal excretion and apparent digestibility of Nellore males with extreme RFI phenotypes (the most and the least efficient terciles) and classified as negative or positive EBV-RFI

| Item | Low<br>RFI | High<br>RFI | SE | P-value | Negative<br>EBV-RFI | Positive<br>EBV-RFI | P-value |
| --- | --- | --- | --- | --- | --- | --- | --- |
| No. of animals | 40 | 40 |  |  | 56 | 24 |  |
| *Performance and feed efficiency traits* |  |  |  |  |  |  |  |
| Initial BW, kg | 256 | 263 | 11.2 | 0.26 | 258±11.4 | 263±12.2 | 0.39 |
| mBW, kg | 298 | 305 | 5.57 | 0.33 | 300±4.82 | 306±7.11 | 0.42 |
| ADG, kg/day | 1.000 | 1.024 | 0.03 | 0.60 | 0.994±0.028 | 1.038±0.043 | 0.39 |
| DMI, kg/day | 6.92 | 8.66 | 0.16 | <0.01 | 7.34±0.15 | 8.82±0.23 | <0.01 |
| DMI, % mBW/day | 2.39 | 2.82 | 0.07 | <0.01 | 2.50±0.06 | 2.86±0.07 | <0.01 |
| FCR | 7.93 | 9.42 | 0.22 | <0.01 | 8.36±0.20 | 9.55±0.30 | <0.01 |
| RFI, kg/day | -0.748 | 0.775 | 0.08 | <0.01 | -0.345±0.093 | 0.833±0.141 | <0.01 |
| EBV-RFI, kg/day | -0.276 | 0.043 | 0.03 | <0.01 | -0.217±0.020 | 0.121±0.029 | <0.01 |
| *Excretion* |  |  |  |  |  |  |  |
| Fecal, kg DM/day | 2.43 | 3.01 | 0.09 | <0.01 | 2.55±0.08 | 3.10±0.12 | <0.01 |
| *Apparent digestibility, g/kg* |  |  |  |  |  |  |  |
| DMD | 661 | 669 | 7.6 | 0.46 | 665±6.42 | 667±9.86 | 0.87 |
| CPD | 607 | 597 | 13.3 | 0.51 | 606±12.5 | 592±16.0 | 0.37 |
| EED | 564 | 573 | 15.8 | 0.64 | 563±13.0 | 586±20.5 | 0.35 |
| NDFD | 461 | 466 | 31.4 | 0.81 | 461±30.2 | 470±33.1 | 0.69 |
| GED | 631 | 634 | 8.5 | 0.75 | 632±7.17 | 631±11.1 | 0.97 |
| NDT | 697 | 699 | 13.2 | 0.87 | 697±12.5 | 700±14.3 | 0.78 |

EBV: estimated breeding value, SE: standard error; mBW<sup>0.75</sup>, midpoint body weight, ADG: average daily gain; DMI: dry matter intake; mBW: midpoint body weight; FCR: feed conversion ratio; RFI: residual feed intake; DMD: dry matter digestibility; CPD: crude protein digestibility; EED: ether extract digestibility; NDFD: neutral detergent fiber digestibility; GED: gross energy digestibility; TDN: total digestible nutrients.

The animals emitted on average 180.2 g CH_4_/day. High RFI animals emitted more 14 g CH_4_/day than low RFI animals. However, low RFI animals (most efficient) emitted more CH_4_ expressed as g/kg DMI (23.1 vs. 20.1 g/day) and lost more gross energy in the form of methane (8.13 vs. 7.08%). There were no differences in methane emission per mBW^0.75^ or ADG. When the animals were classified based on EBV-RFI, no difference in CH_4_ emission (g/day) was observed between most and least efficient animals (Table 3).

**Table 3.** Enteric methane emission of Nellore males with extreme RFI phenotypes (the most efficient tercile and the least efficient tercile) and classified as negative or positive EBV-RFI

| Item | Low<br>RFI | High<br>RFI | SE | P-value | Negative<br>EBV-RFI | Positive<br>EBV-RFI | P-value |
| --- | --- | --- | --- | --- | --- | --- | --- |
| No. of animals | 40 | 40 |  |  | 56 | 24 |  |
| CH<sub>4</sub>, g/day | 174 | 188 | 4.39 | 0.02 | 177±3.70 | 189±5.82 | 0.11 |
| CH<sub>4</sub>, g/kg mBW/day | 0.601 | 0.622 | 0.013 | 0.27 | 0.608±0.011 | 0.621±0.018 | 0.55 |
| CH<sub>4</sub>, g/kg ADG/day | 180 | 186 | 4.98 | 0.39 | 183±4.21 | 185±6.33 | 0.78 |
| CH<sub>4</sub>, g/kg DMI/day | 23.1 | 20.1 | 0.45 | <0.01 | 22.2±0.39 | 19.8±0.62 | <0.01 |
| CH<sub>4</sub>, Mcal/100 Mcal<br>GEI/day | 8.13 | 7.08 | 0.16 | <0.01 | 7.83±0.14 | 6.98±0.22 | <0.01 |

EBV: estimated breeding value; SE: standard error; CH<sub>4</sub>: methane emission expressed in grams per day; mBW: midpoint body weight; ADG: average daily gain; DMI: dry matter intake; GEI: gross energy intake.

The phenotypic correlations between apparent DM and feed compounds digestibility and methane emission are shown in Table 4. Digestibility was not correlated with enteric methane emission, except for EE digestibility which was negatively correlated with CH_4_ (g/day) and CH_4_ (g/kg DM/day), indicating that the higher the EE digestibility, the lower the enteric methane emission.

**Table 4.** Phenotypic correlations between digestibility variables and enteric methane emission

| Item | CH<sub>4</sub>, g/day | CH<sub>4</sub>, g/kg DM/day |
| --- | --- | --- |
| DMD | -0.067<sup>NS</sup> | -0.112<sup>NS</sup> |
| CPD | -0.135<sup>NS</sup> | -0.144<sup>NS</sup> |
| EED | -0.305 (P<0.05) | -0.460 (P<0.01) |
| NDFD | -0.075<sup>NS</sup> | -0.107<sup>NS</sup> |
| GED | -0.058<sup>NS</sup> | -0.102<sup>NS</sup> |
| TDN | -0.075<sup>NS</sup> | -0.112<sup>NS</sup> |

CH<sub>4</sub>, g/day: methane emission expressed as grams per day; CH<sub>4</sub>, g/kg DM/day: methane emission expressed as grams per kg of dry matter intake per day; DMD: dry matter digestibility; CPD: crude protein digestibility; EED: ether extract digestibility; NDFD: neutral detergent fiber digestibility; GED: gross energy digestibility; TDN: total digestible nutrients; NS: non-significant.

## Discussion

There was no difference in the initial body weight, mBW or ADG between animals with extreme RFI phenotypes (the most efficient tercile and the least efficient tercile) despite the large difference in DMI (Table 2). This finding can be explained by the fact that RFI is an efficiency measure that is independent of the growth rate and weight [33]. The average difference in DMI between animals classified as most and least efficient (1.74 kg DM/day) was similar to those reported in other studies involving different animal categories [Fitzsimons et al. [34], 1.91 kg DM/day for Simmental cows; McGee et al. [35], 1.80 kg DM/day for Red Angus steers in the growth phase and 2.40 kg DM/day in the finishing phase]. Various factors might be related to the higher DMI of least efficient animals, including digestibility [5,9]. In general, an increase in DMI reduces diet digestibility because of the increased passage rate of digesta in the rumen; consequently, DM and feed compounds digestibility is expected to be lower in high RFI cattle compared to low RFI animals [15]. In addition, a higher nutrient supply is lost in feces. In a review of 14 studies on the DM digestibility of low versus high RFI animals, Kenny et al. [15] found only one study that reported higher DM digestibility in more efficient animals. The lack of difference in DM digestibility among cattle with varying RFI phenotypes might be related to the type of diet offered since the effect of feed intake on digestion is lower for high-forage diets than for high-concentrate diets. Recent studies have shown higher DM and organic matter digestibility [36] and higher DM and feed compounds digestibility [10] in low RFI animals, even those receiving high-forage diets or high-grain diet [9], and similar digestibility in low and high RFI animals [10]. In indicine cattle, studies have demonstrated higher DM and feed compounds digestibility in low RFI animals compared to high RFI animals fed either a high-forage [8,37] or a high-concentrate diet [38]. On the other hand, Batalha et al. [14] reported lower digestibility for low RFI (most efficient) animals, i.e., a negative relationship between RFI and digestibility. Therefore, the results of the present study showing similar digestibility between animals classified based on extreme phenotypes for RFI and also genetically classified as negative and positive RFI, whose RFI was obtained for a high-forage diet, do not confirm the hypothesis that more efficient animals have higher digestibility, in agreement with the majority of the results reported in the review by Kenny et al. [15].

Low RFI animals excreted −0.581 kg DM/day in feces than high RFI animals (Table 2), an expected result because of the lower DMI [2]. Considering the test period of 83 days, the 40 most efficient animals produced almost two tons (1,929 kg DM) less feces than the least efficient animals, corresponding to 19.3% less excretion and potentially less methane and nitrous oxide release into soil [18]. In addition, this fact reduces concerns regarding waste management [39]. Rius et al. [13] reported lower fecal nitrogen production and higher feed compounds digestibility in lactating cows classified as low RFI.

The benefits of using more efficient animals can be extended to reducing enteric methane emissions while maintaining the same level of production [3,10,17]. Among the factors that influence methane emission, feed intake is the main factor responsible for the variation in daily emissions. Methane emission increases 15 minutes after a feeding event and subsequent fermentation, and this elevated emission can persist for several hours [40]. In beef cattle, CH_4_ (g/day) shows a high phenotypic (0.71) and genetic (0.86) correlation with DMI [41]. If there is a strong positive correlation (phenotypic and genetic) of methane emissions with feed intake, RFI should also show a strong (or at least moderate) positive correlation with methane emission since negative RFI cattle consume less than expected for their live weight and weight gain. It is therefore expected that methane emission will be lower in negative RFI animals proportionally to the lower DMI. Indeed, Manafiazar et al. [4] evaluating a considerable number of animals (heifers and cows) concluded that low RFI animals (from both categories) emit less daily enteric CH_4_ and CO_2_, mainly due to lower feed intake at equal body weight, gain, and fatness. However, the authors also reported that low RFI heifers and cows had higher CH_4_ and CO_2_ per kg of DM than the high RFI counterparts.

In the present study, animals with extreme RFI phenotypes differed in terms of CH_4_ emission (g/day), in which the most efficient animals emitted −7.4% than the least efficient animals (Table 3). Due to the large difference in feed intake between efficiency classes, animals classified as low RFI or as negative EBV-RFI emitted more CH_4_ (g/day) per kg DMI and also expressed as a percentage of gross energy intake lost in the form of methane. No significant difference between RFI classes was found when emission was expressed in relation to ADG or mBW; consequently, most and least efficient animals had similar CH4 expressed in relation to ADG and mBW.

Over the last decade, studies have strongly recommended the use of more efficient animals (low RFI) as an indirect measure to reduce methane emissions from cattle. However, with the increase in the number of studies, this hypothesis did not prove entirely true. The results showed that the association between methane emission and RFI is positive for high-digestibility diets [2,3,6] and that negative RFI animals have lower CH_4_ expressed as g/day and g/kg DM/day. On the other hand, in cattle fed low-digestibility diets such as those that predominate in Brazil, the phenotypic association between methane emission (CH_4_, g/day) and RFI is highly variable (positive, zero or even negative, [3,4,42–44], and that individual enteric methane emission in relation to DMI may be higher in more efficient animals (low RFI). Herd et al. [9] also estimated a negative correlation (−0.54) between CH_4_ expressed in relation to DMI and RFI, even in animals fed a high-digestibility diet.

The associations between the enteric methane emission variables (CH_4_, g/day and CH_4_, g/kg DM/day) and DM and feed compounds digestibility were close to zero, except for EE digestibility (EED, %) which was moderate and negative. The higher EE digestibility and lower methane emission are consistent with the results of Sobrinho et al. [45] who reported a negative correlation between EE intake and methane emission. Likewise, Ellis et al. [46] observed a negative effect of dietary EE content when this variable was included in the regression equation for predicting methane emission. Dietary addition of lipids influences CH_4_ production in ruminants [47] due to the process of biohydrogenation by ruminal microorganisms that adds H2 to the double bonds of unsaturated fatty acids [48], draining the element necessary for the formation of methane [49]. Furthermore, fiber degradation is reduced by the dietary addition of lipids due to the formation a layer that surrounds the fiber and impairs the adhesion of microorganisms [50].

Taken together, the present results demonstrate the economic and environmental benefits of using more efficient animals in the beef production chain, i.e., animals that exhibit lower feed intake, lower fecal excretion and lower enteric methane emission without differences in weight gain or body weight. In cattle farming, feed costs account for the largest part of the production costs and increasing feed efficiency is extremely important because of the reduction in the amount of feed consumed per kilogram of meat produced. Increasing feed efficiency also becomes more important as the production of environmental pollutants such as feces and methane is reduced, meeting the global demands for minimizing the use of inputs and production of waste. However, monitoring the animal fat deposition should be paramount since very lean carcasses have lower market value, and the reduction of feed intake and pollutants may not compensate for the loss value of the final product. The variations in RFI cannot be explained by differences in DM or feed compounds digestibility. More efficient animals emit less enteric methane (CH_4_, g/day) than less efficient animals, probably as a result of lower DMI.

## Supporting information

S1 File. Individual data

## References

1. O’Mara FP. The significance of livestock as a contributor to global greenhouse gas emissions today and in the near future. Anim Feed Sci Technol. 2011;166–167: 7–15. doi:10.1016/j.anifeedsci.2011.04.074

2. Hegarty RS, Goopy JP, Herd RM, McCorkell B. Cattle selected for lower residual feed intake have reduced daily methane production. J Anim Sci. 2007;85: 1479–1486. doi:10.2527/jas.2006-236

3. Jones FM, Phillips FA, Naylor T, Mercer NB. Methane emissions from grazing Angus beef cows selected for divergent residual feed intake. Anim Feed Sci Technol. 2011;166–167: 302–307. doi:10.1016/j.anifeedsci.2011.04.020

4. Manafiazar G, Baron VS, McKeown L, Block H, Ominski K, Plastow G, et al. Methane and carbon dioxide emissions from yearling beef heifers and mature cows classified for residual feed intake under drylot conditions. Can J Anim Sci. 2020;100: 522–535. doi:10.1139/cjas-2019-0032

5. Herd RM, Arthur PF. Physiological basis for residual feed intake. J Anim Sci. 2009;87: E64–E71. doi:10.2527/jas.2008-1345

6. Nkrumah JD, Okine EK, Mathison GW, Schmid K, Li C, Basarab JA, et al. Relationships of feedlot feed efficiency, performance, and feeding behavior with metabolic rate, methane production, and energy partitioning in beef cattle. J Anim Sci. 2006;84: 145–153. doi:10.2527/2006.841145x

7. Krueger W, Carstens G, Gomez R, Bourg B, Lancaster P, Slay L, et al. Relationships between residual feed intake and apparent nutrient digestibility, in vitro methane producing activity and VFA concentrations in growing Brangus heifers. Journal of Animal Science. 2009;87: 129.

8. Oliveira LF, Ruggieri AC, Branco RH, Cota OL, Canesin RC, Costa HJU, et al. Feed efficiency and enteric methane production of Nellore cattle in the feedlot and on pasture. Anim Prod Sci. 2016;58: 886. doi:10.1071/AN16303

9. Herd RM, Velazco JI, Smith H, Arthur PF, Hine B, Oddy H, et al. Genetic variation in residual feed intake is associated with body composition, behavior, rumen, heat production, hematology, and immune competence traits in Angus cattle. J Anim Sci. 2019;97: 2202–2219. doi:10.1093/jas/skz077

10. Johnson JR, Carstens GE, Krueger WK, Lancaster PA, Brown EG, Tedeschi LO, et al. Associations between residual feed intake and apparent nutrient digestibility, in vitro methane-producing activity, and volatile fatty acid concentrations in growing beef cattle. J Anim Sci. 2019;97: 3550–3561. doi:10.1093/jas/skz195

11. Lawrence P, Kenny DA, Earley B, McGee M. Grazed grass herbage intake and performance of beef heifers with predetermined phenotypic residual feed intake classification. animal. 2012;6: 1648–1661. doi:10.1017/S1751731112000559

12. Lawrence P, Kenny DA, Earley B, Crews DH, McGee M. Grass silage intake, rumen and blood variables, ultrasonic and body measurements, feeding behavior, and activity in pregnant beef heifers differing in phenotypic residual feed intake1. J Anim Sci. 2011;89: 3248–3261. doi:10.2527/jas.2010-3774

13. Rius AG, Kittelmann S, Macdonald KA, Waghorn GC, Janssen PH, Sikkema E. Nitrogen metabolism and rumen microbial enumeration in lactating cows with divergent residual feed intake fed high-digestibility pasture. J Dairy Sci. 2012;95: 5024–5034. doi:10.3168/jds.2012-5392

14. Batalha CDA, Morelli M, Branco RH, Cyrillo JNSG, Canesin RC, Mercadante MEZ, et al. Association between residual feed intake, digestion, ingestive behavior, enteric methane emission and nitrogen metabolism in Nellore beef cattle. Animal Science Journal. 2020;91. doi:10.1111/asj.13455

15. Kenny DA, Fitzsimons C, Waters SM, McGee M. Invited review: Improving feed efficiency of beef cattle – the current state of the art and future challenges. Animal. 2018;12: 1815–1826. doi:10.1017/S1751731118000976

16. Subepang S, Suzuki T, Phonbumrung T, Sommart K. Enteric methane emissions, energy partitioning, and energetic efficiency of zebu beef cattle fed total mixed ration silage. Asian-Australas J Anim Sci. 2019. doi:10.5713/ajas.18.0433

17. Fitzsimons C, Kenny DA, Deighton MH, Fahey AG, McGee M. Methane emissions, body composition, and rumen fermentation traits of beef heifers differing in residual feed intake. J Anim Sci. 2013;91: 5789–5800. doi:10.2527/jas.2013-6956

18. Herd R, Arthur P, Hegarty R. Potential to reduce greenhouse gas emissions from beef production by selection to reduce residual feed intake. 7th World Congr. Genet. Anim. Prod. Montpellier, France; 2002. pp. 10–22.

19. Cantalapiedra-Hijar G, Abo-Ismail M, Carstens GE, Guan LL, Hegarty R, Kenny DA, et al. Review: Biological determinants of between-animal variation in feed efficiency of growing beef cattle. Animal. 2018;12: s321–s335. doi:10.1017/S1751731118001489

20. Benfica LF, Sakamoto LS, Magalhães AFB, de Oliveira MHV, de Albuquerque LG, Cavalheiro R, et al. Genetic association among feeding behavior, feed efficiency, and growth traits in growing indicine cattle. J Anim Sci. 2020;98. doi:10.1093/jas/skaa350

21. NRC. Nutrient Requirements of Beef Cattle. 7th ed. Washington, D.C.: National Academy Press; 2000.

22. Sampaio CB, Detmann E, Valente TNP, Costa VAC, Valadares Filho S de C, Queiroz AC de. Fecal excretion patterns and short term bias of internal and external markers in a digestion assay with cattle. Revista Brasileira de Zootecnia. 2011;40: 657–665. doi:10.1590/S1516-35982011000300026

23. Casali AO, Detmann E, Valadares Filho S de C, Pereira JC, Henriques LT, Freitas SG de, et al. Influência do tempo de incubação e do tamanho de partículas sobre os teores de compostos indigestíveis em alimentos e fezes bovinas obtidos por procedimentos in situ. Revista Brasileira de Zootecnia. 2008;37: 335–342. doi:10.1590/S1516-35982008000200021

24. Mertens DR. Gravimetric Determination of Amylase-Treated Neutral Detergent Fiber in Feeds with Refluxing in Beakers or Crucibles: Collaborative Study. J AOAC Int. 2002;85: 1217–1240.

25. Detmann E, Paulino MF, Zervoudakis JT, Valadares Filho S de C, Euclydes RF, Lana R de P, et al. Cromo e indicadores internos na determinação do consumo de novilhos mestiços, suplementados, a pasto. Revista Brasileira de Zootecnia. 2001;30: 1600–1609. doi:10.1590/S1516-35982001000600030

26. Cochran RC, Galyean ML. Measurement of in vivo Forage Digestion by Ruminants. In: George C. Fahey Jr., editor. Forage Quality, Evaluation, and Utilization. American Society of Agronomy; 1994. pp. 613–643. doi:10.2134/1994.foragequality.c15

27. Berndt A, Boland TB, Deighton M, Gere J, Grainger C, Hegarty R, et al. Guidelines for use of sulphur hexafluoride (SF6) tracer technique to measure enteric methane emissions from ruminants. Jonke A, Waghorn G, editors. Wellington, NZ: Ministry for Primary Industries; 2014.

28. Deighton MH, Williams SRO, Hannah MC, Eckard RJ, Boland TM, Wales WJ, et al. A modified sulphur hexafluoride tracer technique enables accurate determination of enteric methane emissions from ruminants. Anim Feed Sci Technol. 2014;197: 47–63. doi:10.1016/j.anifeedsci.2014.08.003

29. AOAC. Official Methods of Analytical of AOAC International. 15th ed. Washington, DC: Assoc. Off. Anal. Chem.; 1990.

30. AOAC International. Official Methods of Analysis. 18th ed. Arlington, VA: AOAC International; 2005.

31. van Soest PJ, Robertson JB, Lewis BA. Methods for Dietary Fiber, Neutral Detergent Fiber, and Nonstarch Polysaccharides in Relation to Animal Nutrition. J Dairy Sci. 1991;74: 3583–3597. doi:10.3168/jds.S0022-0302(91)78551-2

32. Parr Instrument Company. Oxygen bomb calorimetry and combustion methods. Moline, IL: Parr Instrument Co; 1960.

33. Koch RM, Swiger LA, Chambers D, Gregory KE. Efficiency of Feed Use in Beef Cattle. J Anim Sci. 1963;22: 486–494. doi:10.2527/jas1963.222486x

34. Fitzsimons C, Kenny DA, Fahey AG, McGee M. Feeding behavior, ruminal fermentation, and performance of pregnant beef cows differing in phenotypic residual feed intake offered grass silage. J Anim Sci. 2014;92: 2170–2181. doi:10.2527/jas.2013-7438

35. McGee M, Welch CM, Ramirez JA, Carstens GE, Price WJ, Hall JB, et al. Relationships of feeding behaviors with average daily gain, dry matter intake, and residual feed intake in Red Angus–sired cattle. J Anim Sci. 2014;92: 5214–5221. doi:10.2527/jas.2014-8036

36. de La Torre A, Andueza D, Renand G, Baumont R, Cantalapiedra-Hijar G, Nozière P. Digestibility contributes to between-animal variation in feed efficiency in beef cows. Animal. 2019;13: 2821–2829. doi:10.1017/S1751731119001137

37. Magnani E, Nascimento CF, Branco RH, Bonilha SFM, Ribeiro EG, Mercadante MEZ. Relações entre consumo alimentar residual, comportamento ingestivo e digestibilidade em novilhas Nelore. Boletim de Indústria Animal. 2013;70: 187–194. doi:10.17523/bia.v70n2p187

38. Bonilha SFM, Branco RH, Mercadante MEZ, dos Santos Gonçalves Cyrillo JN, Monteiro FM, Ribeiro EG. Digestion and metabolism of low and high residual feed intake Nellore bulls. Trop Anim Health Prod. 2017;49: 529–535. doi:10.1007/s11250-017-1224-9

39. Montes F, Meinen R, Dell C, Rotz A, Hristov AN, Oh J, et al. SPECIAL TOPICS — Mitigation of methane and nitrous oxide emissions from animal operations: II. A review of manure management mitigation options. J Anim Sci. 2013;91: 5070–5094. doi:10.2527/jas.2013-6584

40. de Haas Y, Pszczola M, Soyeurt H, Wall E, Lassen J. Invited review: Phenotypes to genetically reduce greenhouse gas emissions in dairying. J Dairy Sci. 2017;100: 855–870. doi:10.3168/jds.2016-11246

41. Donoghue KA, Bird-Gardiner T, Arthur PF, Herd RM, Hegarty RF. Genetic and phenotypic variance and covariance components for methane emission and postweaning traits in Angus cattle. J Anim Sci. 2016;94: 1438–1445. doi:10.2527/jas.2015-0065

42. Freetly HC, Brown-Brandl TM. Enteric methane production from beef cattle that vary in feed efficiency. J Anim Sci. 2013;91: 4826–4831. doi:10.2527/jas.2011-4781

43. Mercadante MEZ, Caliman AP de M, Canesin RC, Bonilha SFM, Berndt A, Frighetto RTS, et al. Relationship between residual feed intake and enteric methane emission in Nellore cattle. Revista Brasileira de Zootecnia. 2015;44: 255–262. doi:10.1590/S1806-92902015000700004

44. Velazco JI, Herd RM, Cottle DJ, Hegarty RS. Daily methane emissions and emission intensity of grazing beef cattle genetically divergent for residual feed intake. Anim Prod Sci. 2016;57: 627. doi:10.1071/AN15111

45. Sobrinho TLP, Branco RH, Magnani E, Berndt A, Canesin RC, Mercadante MEZ. Development and evaluation of prediction equations for methane emission from Nellore cattle Dry matter intake (DMI). Acta Sci. 2018;41: 42559. doi:10.4025/actascianimsci.v41i1.42559

46. Ellis JL, Kebreab E, Odongo NE, McBride BW, Okine EK, France J. Prediction of Methane Production from Dairy and Beef Cattle. J Dairy Sci. 2007;90: 3456–3466. doi:10.3168/jds.2006-675

47. Beauchemin KA, Kreuzer M, O’Mara F, McAllister TA. Nutritional management for enteric methane abatement: a review. Aust J Exp Agric. 2008;48: 21. doi:10.1071/EA07199

48. Machmüller A, Kreuzer M. Methane suppression by coconut oil and associated effects on nutrient and energy balance in sheep. Can J Anim Sci. 1999;79: 65–72. doi:10.4141/A98-079

49. Jenkins TC. Lipid Metabolism in the Rumen. J Dairy Sci. 1993;76: 3851–3863. doi:10.3168/jds.S0022-0302(93)77727-9

50. Valadares Filho S, Pina D. Fermentação ruminal. In: Berchielli T, Pires A, Oliveira S, editors. Nutrição de Ruminantes. 2006. pp. 151–182.

